# Platelet-programmed circulating tumor cells signal to monocytes through a candidate CD40LG–ITGA5:ITGB1 myeloid checkpoint axis in breast cancer

**DOI:** 10.64898/2026.09.20.752972

**Authors:** Samane Khoshbakht, Hojat Borna, Mehrnaz Zarei, Zahra Salehi, Niloofar Hejazifar, Yasaman Setayeshpour, Niroshana Anandasabapathy, Mayte Suarez-Farinas

## Abstract

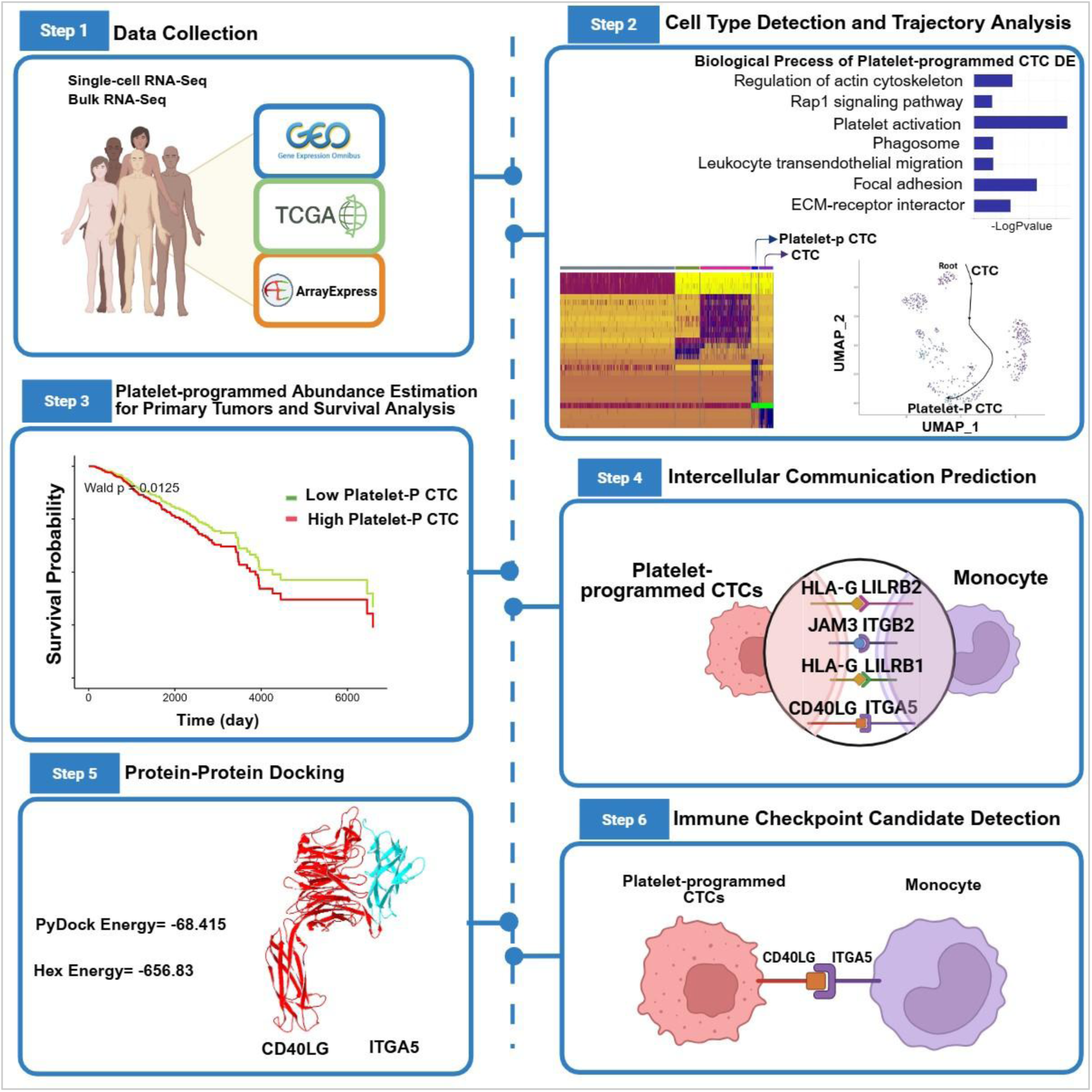

Circulating tumor cells (CTCs) are key drivers of distant metastasis, while platelets facilitate this process by protecting them from immune surveillance. However, the detection of CTCs exhibiting platelet markers and their communication with monocytes has not been thoroughly investigated.

This study seeks to identify the intercellular communication of CTCs that have acquired platelet traits by integrating single-cell RNA-seq data of 377 CTCs and 2,634 white blood cells. Trajectory analysis identified a subpopulation of CTCs with enhanced platelet-related functions, including platelet aggregation and resistance to NK cell–mediated cytotoxicity, termed Platelet-Programmed CTCs. The Dampened Weighted Least Squares method was implemented to estimate the platelet-programmed tumor cell proportion in 1,102 bulk RNA-seq samples from primary breast cancer tumors. A strong inverse association was found between platelet-programmed tumor cell proportion and overall survival (Cox-PH p-value =0.00419, HR=6 per 10% increase in proportion, 95% CI:1.76-20.7). However, this association did not differ significantly between early- and late-stage patients. Cell-cell communication analyses and molecular docking revealed the CD40LG– ITGA5:ITGB1 pair as a potential immune checkpoint candidate for monocytes in cancer (Hex score=−656.83; TNFα–TNFR reference complex=−515). However, further experimental and clinical validation is required to translate these findings. In conclusion, these findings deepen our understanding of the immune evasion mechanisms of CTCs by acquiring platelet characteristics. This immune checkpoint axis offers a novel avenue for further experimental validation and future intervention.

## 1 INTRODUCTION

Circulating tumor cells (CTCs) are cancer cells that detach from the primary tumor and enter the bloodstream, where they face immune surveillance [1]. Beyond their role in dissemination, CTCs have emerged as valuable tools for assessing tumor heterogeneity, informing biomarker-driven immunotherapies, and capturing diverse mechanisms of drug resistance, as well as for detecting minimal residual disease. As such, CTCs provide a unique window into tumor biology at the single-cell level and complement minimally invasive liquid biopsy approaches, with the potential to uncover previously unrecognized immune checkpoint interactions [2].

To persist in the bloodstream, CTCs may acquire durable transcriptional adaptations, potentially shaped by direct or indirect interactions with platelets [3]. Here, we refer to such CTCs with platelet traits as platelet-programmed CTCs (Platelet-P CTCs). Despite evidence that platelet-associated phenotypes support CTC survival, their influence on monocyte immune checkpoint pathways remains undefined, limiting our understanding of immune evasion in circulation.

Monocyte-associated immune checkpoint pathways, defined by inhibitory receptor–ligand interactions within the tumor microenvironment, are increasingly recognized as important regulators of anti-tumor immunity[4]. Growing evidence identifies these myeloid checkpoints as a determinant of therapeutic response, contributing to tumor-driven immune dysfunction [5]. Monocyte-associated immune checkpoint pathways represent a critical but underexplored axis of tumor immune evasion in the circulation. Two key suppressive axes have been identified: the HLA-G–LILRB1/LILRB2 pathway, through which tumor-derived HLA-G engages inhibitory LILRB receptors on monocytes to promote immune tolerance and facilitate escape from cytotoxic immunity; and the PD-1–PD-L1 axis, whereby PD-L1 expression on circulating monocytes suppresses T-cell activity and has been shown to correlate with disease progression and tumor burden in advanced breast cancer [5–9]. Together, these pathways position monocytes not only as mediators of immune suppression but also as accessible biomarkers of systemic tumor activity [6, 9]. Despite this, it remains unknown how these checkpoint pathways are regulated in the circulation—and specifically, whether Platelet-P CTCs actively exploit monocyte checkpoint signaling to evade immune clearance. Addressing this gap may reveal previously unrecognized mechanisms of immune suppression and identify novel checkpoint interactions with therapeutic potential.

Given these observations, we sought to determine whether Platelet-P CTCs modulate immune checkpoint signaling in monocytes through intercellular communication networks. By integrating publicly available single-cell RNA sequencing datasets, we identified CD40LG–ITGA5 as a candidate immunoregulatory ligand–receptor interaction potentially mediating CTC–monocyte crosstalk. Notably, a higher platelet-programmed tumor cell proportion was significantly associated with poorer overall survival in patients with breast cancer.

## 2 METHODS AND MATERIALS

### 2.1 Data Query

Single-cell RNA-seq datasets of circulating tumor cells (CTCs) and white blood cells (WBCs) were collected from the NCBI and EBI repositories. Our selection criteria excluded data from cell lines and solid tumor samples (both primary and metastatic), retaining only those that profiled single CTCs and CD45⁺ WBCs isolated from patient blood. Cell identity was taken from the annotations of the original studies rather than re-derived here: only cells annotated by the source authors as single CTCs were retained, whereas cells annotated as CTC clusters or CTC–white blood cell clusters were excluded. The consort diagram is presented in Fig.1. An initial search of the NCBI GEO database using “circulating tumor cells” [All Fields] AND single-cell [All Fields] AND “Homo sapiens” [Organism] retrieved 134 records. A subsequent query, “circulating tumor cells” [All Fields] OR ctc [All Fields] AND (“Homo sapiens” [Organism] AND “Expression profiling by high throughput sequencing” [Filter] AND “attribute name tissue” [Filter]), yielded 36 records. An additional search in EBI returned overlapping entries.

**Fig 1.**
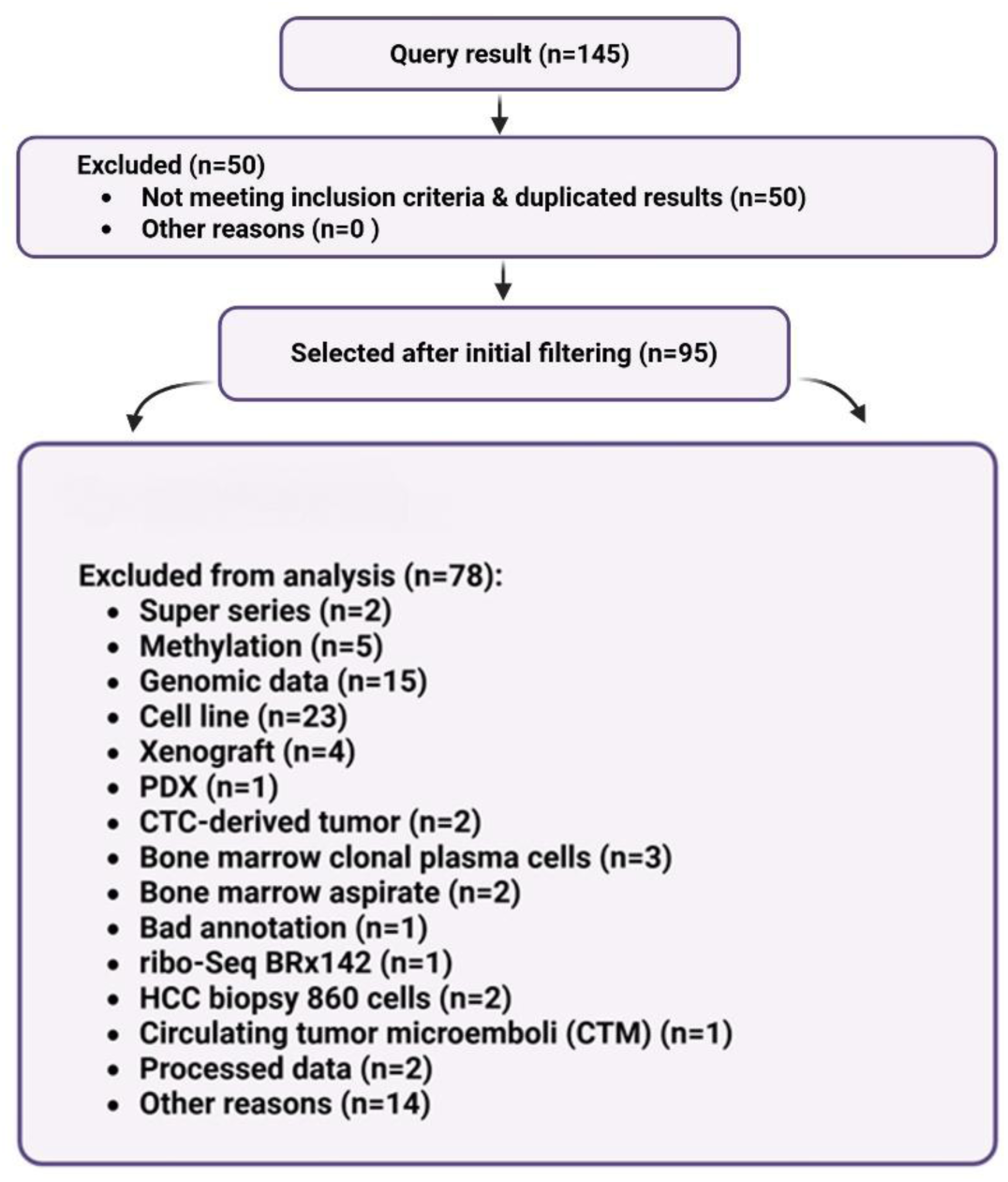
CONSORT diagram for the studies contributing to this analysis. “n” refers to the unique number of single-cell experiments with unique GEO numbers.

To support our cell deconvolution analysis, we analyzed 1,102 bulk mRNA gene expression profiles from primary breast cancer tumors available in the Cancer Genome Atlas (TCGA) database. Based on the AJCC staging information provided in the accompanying metadata, patients with stage I or II disease were classified as early-stage, while those with stage III or IV disease were categorized as late-stage (Supplementary File 1).

### 2.2 Normalization and Integration Within-study preprocessing

For each study, the count matrix of CTCs/WBCs was preprocessed, gene symbols were annotated using Biomart, and individual Seurat objects were created in R. The metadata of each Seurat object was added separately. CTCs with fewer than 2,500 or more than 100,000 mRNA counts were filtered out, so that only cells with a reliable transcriptome were retained; for WBCs, cells with at least 2,500 detected genes and at most 25% mitochondrial reads were retained. All CTC datasets were generated by full-length single-cell RNA sequencing of individually isolated cells, each processed in a separate reaction (micromanipulation into individual tubes or wells for GSE75367, GSE109761, GSE111065, GSE144494, GSE144495 and GSE180097; single-cell microfluidic chambers for GSE186288) using template-switching chemistry, rather than droplet-based capture [10–15]. Ambient-RNA contamination arises from cell-free mRNA co-encapsulated with cells in a shared droplet emulsion and therefore cannot occur in these libraries; accordingly, ambient-RNA correction tools (e.g., SoupX) were not applied to the CTC data [16]. The white blood cell dataset (GSE174461) was generated on the droplet-based 10x Chromium platform from red-blood-cell-lysed, FACS-sorted, viability-checked PBMCs [17], a preparation that limits the cell-free mRNA contributing to ambient contamination. No dedicated ambient-RNA correction was applied to this dataset; instead, low-content barcodes, in which ambient contamination is proportionally greatest, were removed by the stringent cell-level filtering described above, and receptor-level analyses in monocytes were restricted to genes detected in at least 25% of cells, well above the few percent of counts typically attributable to ambient RNA[16].

#### Cross-study Integration

To integrate the datasets, we applied Seurat’s anchor-based integration workflow [18]. Before integration, each dataset was independently normalized using SCTransform to model study-specific technical variation, followed by anchor-based integration across datasets. Integration was applied to harmonize cell-state relationships across studies, rather than to enable direct comparison of gene expression magnitudes. Feature selection was performed using the variance stabilizing transformation method, selecting the top 2,000 highly variable genes. For dimensionality reduction, principal component analysis (PCA) was used to capture the major sources of variation, and the 10 top principal components were subsequently used to generate a two-dimensional embedding via Uniform Manifold Approximation and Projection (UMAP) for visualization and clustering [18, 19]. To identify CTC subtypes, we used an unsupervised k-nearest neighbor (KNN) algorithm with Euclidean distances in PCA space. This function builds a Shared Nearest Neighbor (SNN) graph for the gene expression data. Using this KNN graph, the SNN graph is created by computing the neighborhood overlap between each cell and its k closest neighbors. Clustering was then performed with modularity optimization (resolution parameter =3). To identify differentially expressed genes (DEGs) among cell types, we applied a likelihood-ratio test using a one-vs-rest approach, with the minimum percentage of expressing cells set to 0.25 and a log2 fold-change threshold of 0.25. DEGs for the Platelet-P CTC group were functionally enriched using the EnrichR (FDR < 0.05) [20]. P-values were adjusted using the Benjamini-Hochberg approach, which controls the FDR.

### 2.3 Annotation

We annotated the detected CTC and WBC subpopulations using marker genes and different computational methods, including BlueprintEncodeData, MonacoImmuneData, Human Primary Cell Atlas (HPCA), Database of Immune Cell Expression (DICE), and marker genes [21–23].

### 2.4 CTC to Platelet-Programmed CTC Development Trajectory

To trace the developmental path from CTCs to Platelet-P CTCs and extract the developmentally activated pathways, we applied the Slingshot technique to both CTC and Platelet-Programmed CTC clusters following re-clustering, and dimensionality reduction via UMAP. In this method, we identified the global trajectory structure with a cluster-based minimum spanning tree and fitted the simultaneous principal curves to describe the development [24]. After the construction of the trajectory curve, we fitted a generalized additive model for each gene to model the potentially nonlinear relationships between gene expression and pseudo-time. The root of the trajectory was selected based on entropy analysis. We then detected temporally dynamic genes [24]. Finally, the biological pathways of significant genes that were functionally enriched were analyzed with EnrichR (FDR < 0.05)[20].

### 2.5 Intercellular Communication Process

We investigated the intercellular communication between ligands and receptors for specific cell groups using the NicheNet method [25]. In this method, we had to select the sender (ligands) and receiver (receptors) cell clusters, as well as target genes. To do this, we selected the Platelet-Programmed CTC cluster as the sender group and monocytes as the receiver cells. Therefore, the ligands were detected on Platelet-P CTCs, and receptors and target genes were detected on monocytes.

The original NicheNet method ranks ligands based only on ligand activity [25]. Ligand activity in this method is calculated by comparing the predicted target genes of each ligand, derived from a prior weighted network of ligand–receptor, signaling, and transcriptional interactions, with the observed differentially expressed genes in receiver cells. For each ligand, a regulatory potential score is computed for all possible target genes using network propagation, and ligand activity is quantified based on how well these predicted targets overlap with and rank the observed gene expression changes, typically using metrics such as the area under the precision–recall curve (AUPR). Here, we ranked the ligands not only using the ligand activity but also according to the upregulation of ligand/receptor compared to other cell types. The prioritization weights were selected as the default. Putative receptors for ligands were detected by querying all ligand-receptor databases of NicheNet sources. As the background, all other genes were considered to be expressed in the receiver group [25]. NicheNet utilizes human gene expression data from interacting cells and integrates it with a pre-existing model, incorporating known information on signaling pathways from ligands to targets. This fusion enables the prediction of ligand-receptor interactions that could potentially influence gene expression alterations in specific cells. Finally, to assess the potential of the predicted ligand–receptor pairs as immune checkpoints, we conducted a targeted literature review focusing on the top-ranked interactions.

### 2.6 Tumor Cell Proportion Estimation and Association to Survival

Using the dampened weighted least squares (DWLS) method [26], we estimated the proportion of tumor cells expressing the platelet-programmed transcriptional program—the signature derived from Platelet-P CTCs—in bulk RNA-seq data from 1,102 TCGA primary breast tumors; hereafter, platelet-programmed tumor cell proportion. The signature matrix was identified using the Model-based Analysis of Single-cell Transcriptomics (MAST), applying a hurdle model. Genes were considered markers if they exhibited a false discovery rate (FDR)–adjusted p-value < 0.01 and a log2 fold change greater than 0.5 when comparing the normal and cancer groups. Single-cell transcriptomics reveal gene expression heterogeneity, but are challenged by technical dropouts and bimodal expression patterns, where genes are either highly expressed or undetectable; MAST addresses these issues [27]. To assess the prognostic impact of platelet-programmed tumor cell proportion, multivariable Cox proportional hazards (Cox-PH) regression models were fitted, adjusting for age at diagnosis, PAM50 molecular subtype, and stage (early and late). The estimated platelet-programmed tumor cell proportion was modeled both continuously and dichotomized at the median [28].

### 2.7 Molecular Docking of Ligand-Receptor Pairs

To study the strength of interaction between the predicted immune checkpoint candidate at the protein level, the NMR or X-ray crystallographic structure of the proteins was obtained from the Protein Data Bank (PDB). 3D structures of the proteins were cleaned with Swiss-PdbViewer and UCSF Chimera [29]. The needed subunits were selected and saved in separate files for docking procedures. Two different online and offline docking approaches were utilized. Rigid protein-protein docking algorithms were chosen to be utilized, which are mostly based on shape complementarity, electrostatic interactions, and Van der Waals forces.

PyDock was used as an initial evaluation step for docking [30]. Default settings and no restraints were used for the docking process. However, to increase the rigor of the docking results, protein pairs were also docked with Hex software [30]. To accelerate the calculation process of shape complementarity and electrostatic interactions, Hex uses spherical polar Fourier (SPF) correlations. Both approaches give the total interaction energy. This energy is completely relative to and different among protein pairs. As protein structure grows, the minimum interaction energy to stabilize the protein-protein structure increases.

### 2.8 Annotation of ITGB1 Downstream Signalling in Monocytes

Because ITGA5 was predicted as the monocyte receptor for CD40LG and functions as an obligate heterodimer with ITGB1, we assembled a targeted panel of downstream effectors of ITGA5:ITGB1 (α5β1) signalling through a structured literature review. Genes were assigned a priori to four functional tiers: proximal adhesion signalling (PTK2B, TLN1, VCL), intermediate transcription factors (NFKB2, RELB, REL), cytokine and cytokine-receptor genes (IL10RA, CSF1R, CXCL8, HAVCR2), and tumour-associated macrophage (TAM)-associated signalling programmes (TGFBR1, TGFBR2, TGFBI, S100A9, VEGFA). Expression of this panel was then examined across the annotated CTC and WBC clusters and visualised as a z-scored heatmap. The functional group labels shown in Fig.7b were derived from EnrichR pathway annotation of this curated gene set against the Reactome 2024 database; because the panel was itself selected based on prior pathway membership, the enrichment output was used only to organise genes into functional categories and not as an independent statistical test of pathway activation. The full enrichment output is provided in Supplementary File 3.

## 3 RESULTS

### 3.1 Data cleaning

After QC assessment, gene expression profiles from 8 out of 17 CTC and WBC datasets were selected for the final downstream analyses (Table 1, Fig.1). After removing circulating tumor cells that did not express or showed low expression levels of CTC marker genes, along with those with minimal or no expression of white blood cell marker genes (Supplementary Table.1, Supplementary Fig.1, 2), we retained 377 CTCs out of 706 and 2,634 WBCs.

**Table 1.** CTC and WBC data information.

| Datasets | Cancer | Cell type | n-WBCs | n-CTCs | n-Patients | Age Range | Library method |
| --- | --- | --- | --- | --- | --- | --- | --- |
| GSE186288 | Breast | CTC |  | 81 | 5 | 62-75 | Polaris IFC, full-length (template-switching) |
| GSE180097 | Breast | CTC |  | 36 | 1 | 58 | Smart-seq2, micromanipulation |
| GSE144495 | Breast | CTC |  | 195 | 6 | 62-74 | SMART-seq, CTC-iChip + micromanipulation |
| GSE144494 | Breast | CTC |  | 135 | 9 | 63-74 | SMART-seq, CTC-iChip + micromanipulation |
| GSE109761 | Breast | CTC |  | 116 | 13 | 42-75 | Smart-seq2, Parsortix + micromanipulation |
| GSE111065 | Breast | CTC |  | 69 | 18 | 50-76 | Smart-seq2, Parsortix + micromanipulation |
| GSE75367 | Breast | CTC |  | 74 | 13 | 54-75 | SMARTer, CTC-iChip + micromanipulation |
| GSE174461 | Breast | WBC | 2,634 | 0 | 3 | 59-75 | 10x Chromium |
| TOTAL |  |  | 2,634 | 706 | 68 | 42-75 |  |

Table 1 CTC and WBC data information
| Datasets | Cancer | Cell type | n-WBCs | n-CTCs | n-Patients | Age Range | Library method |
| --- | --- | --- | --- | --- | --- | --- | --- |
| GSE186288 | Breast | CTC |  | 81 | 5 | 62-75 | Polaris IFC, full-length (template-switching) |
| GSE180097 | Breast | CTC |  | 36 | 1 | 58 | Smart-seq2, micromanipulation |
| GSE144495 | Breast | CTC |  | 195 | 6 | 62-74 | SMART-seq, CTC-iChip + micromanipulation |
| GSE144494 | Breast | CTC |  | 135 | 9 | 63-74 | SMART-seq, CTC-iChip + micromanipulation |
| GSE109761 | Breast | CTC |  | 116 | 13 | 42-75 | Smart-seq2,<br>Parsortix +<br>micromanipulation |
| GSE111065 | Breast | CTC |  | 69 | 18 | 50-76 | Smart-seq2,<br>Parsortix +<br>micromanipulation |
| GSE75367 | Breast | CTC |  | 74 | 13 | 54-75 | SMARTer, CTC-iChip +<br>micromanipulation |
| GSE174461 | Breast | WBC | 2,634 | 0 | 3 | 59-75 | 10x Chromium |
| TOTAL |  |  | 2,634 | 706 | 68 | 42-75 |  |

### 3.2 CTC/WBC Subpopulation Annotation

The Platelet-Programmed CTC group was annotated based on elevated expression of canonical platelet markers (n=214), including SELP, PPBP, and RGS18, and confirmed by computational methods (Fig.2a and b, Supplementary Fig.3, Supplementary Table.1). The second CTC cluster (n=163) exhibited CTC markers, such as KRT8 or EPCAM, which are canonical CTC markers (Fig.2a and b, Supplementary Table.1). Moreover, we annotated the other three clusters for WBCs comprising T cells (n=342), NK/NK-T cells (n=704), and monocytes (n=1,588), which are shown as olive green, magenta, and dark grey, respectively (Fig.2a,b, Supplementary Fig.3). These three clusters were PTPRC-positive (CD45-positive) (Fig.2a and b). Given that CD3E expression was observed in NK cells, alongside NK markers like FCGR3A, KLRD1, and PRF1, we labeled this cluster as NK/NK-T cells (magenta cluster in Fig.2b). Monocytes were identified based on gene expression patterns, particularly genes like CD14 and VCAN, while T cells were annotated based on the expression of genes, such as CD3E or CD3D (dark grey and olive green clusters in Fig.2b). Additionally, the platelet-programmed CTC cluster showed differential gene expression enriched in biological processes related to platelet activation and focal adhesion (Fig.2c).

**Fig 2.**
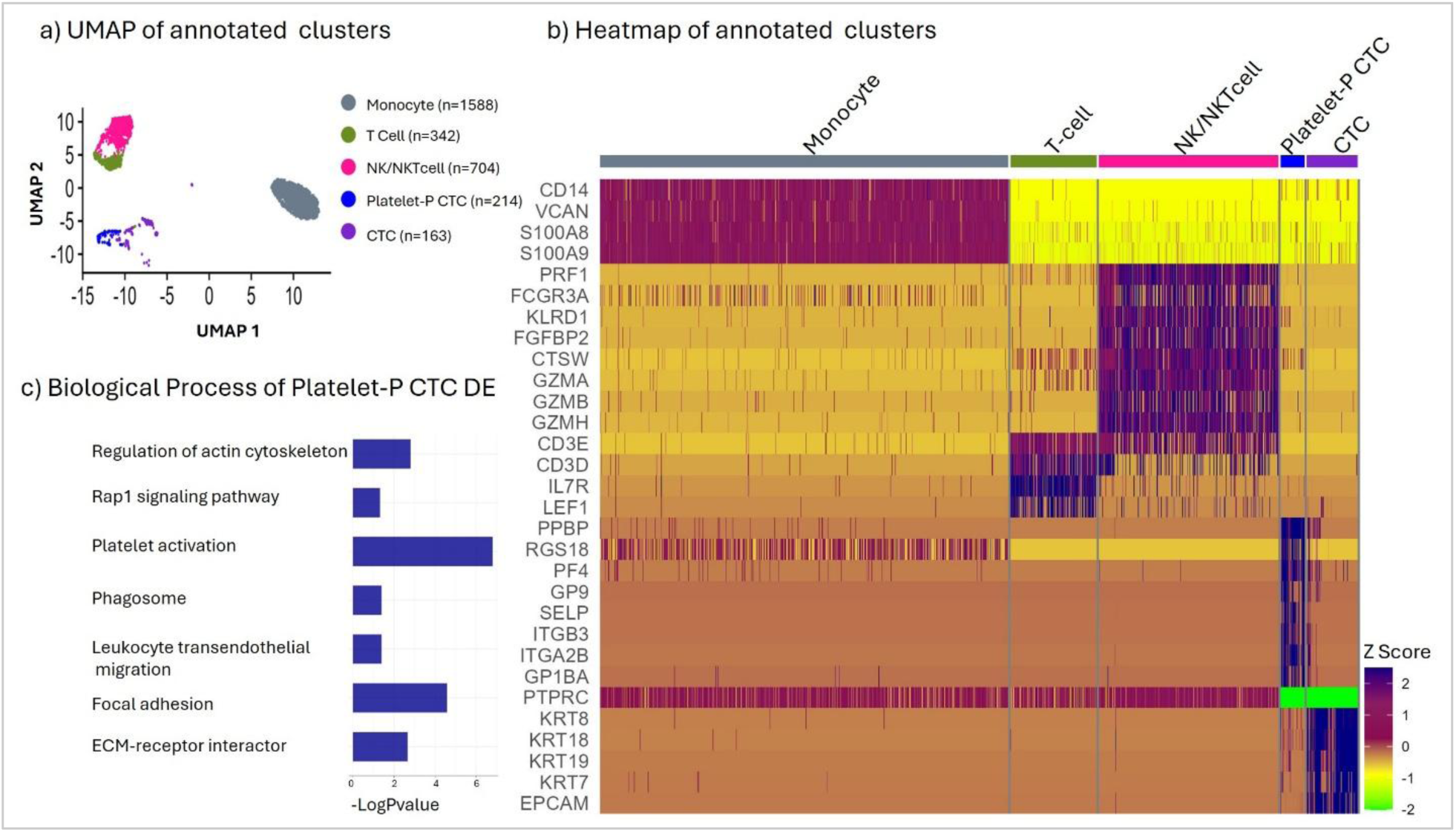
UMAP and Heatmap of detected clusters. a) Each cell type was illustrated by different colors on the UMAP plot. b) Each cluster on the Heatmap plot was annotated with a unique color. The gene list on the left side of the heatmap is the cell-type marker genes. c) Biological processes of Platelet-Programmed CTCs were illustrated by a bar chart. Platelet-P CTCs: Platelet-Programmed CTCs

### 3.3 From CTCs to Platelet-Programmed CTCs: A Progressive Transcriptional Transition

Reclustering of CTC-related clusters and UMAP visualization revealed a continuous transcriptional trajectory from conventional CTCs toward the Platelet-Programmed CTC population, consistent with a progressive acquisition of platelet-associated gene expression programs rather than a discrete phenotypic switch (Fig.3a and b). Along this trajectory, gene sets governing platelet aggregation, homotypic cell-cell adhesion, and protection from NK cell-mediated cytotoxicity were progressively enriched, suggesting that Platelet-P CTCs undergo coordinated transcriptional reprogramming that simultaneously enhances intercellular adhesion and confers immune evasion capacity (Fig.3c,FDR<0.05)

**Fig 3.**
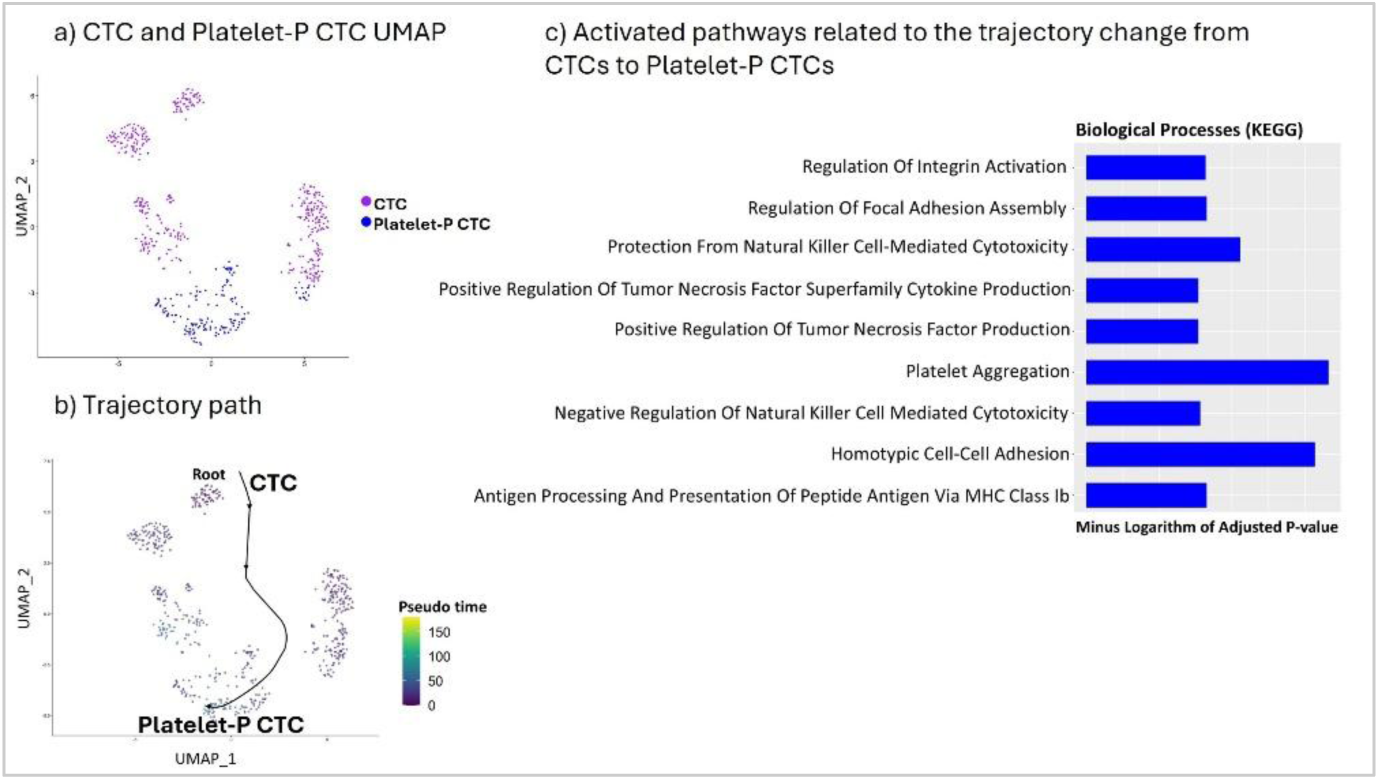
Trajectory path from CTCs to Platelet-programmed CTCs. a) The purple and blue dots in the UMAP plot indicate the CTC and Platelet-Programmed CTC clusters. b) The curve indicates the trajectory path for CTCs and Platelet-Programmed CTC cells. c) The length of bars indicates the level of significance of biological processes for trajectory development. Platelet-P CTC: Platelet-Programmed CTC

### 3.4 Elevated Platelet-Programmed Tumor Cell Proportion is Associated with Shorter OS in Breast Cancer Patients

In the Cox-PH model with the platelet-programmed tumor cell proportion dichotomised at the cohort median, patients in the low-proportion group had significantly longer overall survival than those in the high-proportion group (HR=1.5, p-value=0.012, 95% CI:1.1-2.2) (Fig.4). The same inverse association was observed when the proportion was modelled continuously (p-value=0.004, HR=6, 95% CI:1.76-20.7, Supplementary File 1). Neither stage nor PAM50 subtype modified the association (stage-pvalue= 0.7, subtype-pvalue=0.9). Stratified estimates were directionally consistent: per 10% increase in platelet-programmed tumor cell proportion, the hazard ratio was 1.9 in early-stage and 1.5 in late-stage patients.

**Fig 4.**
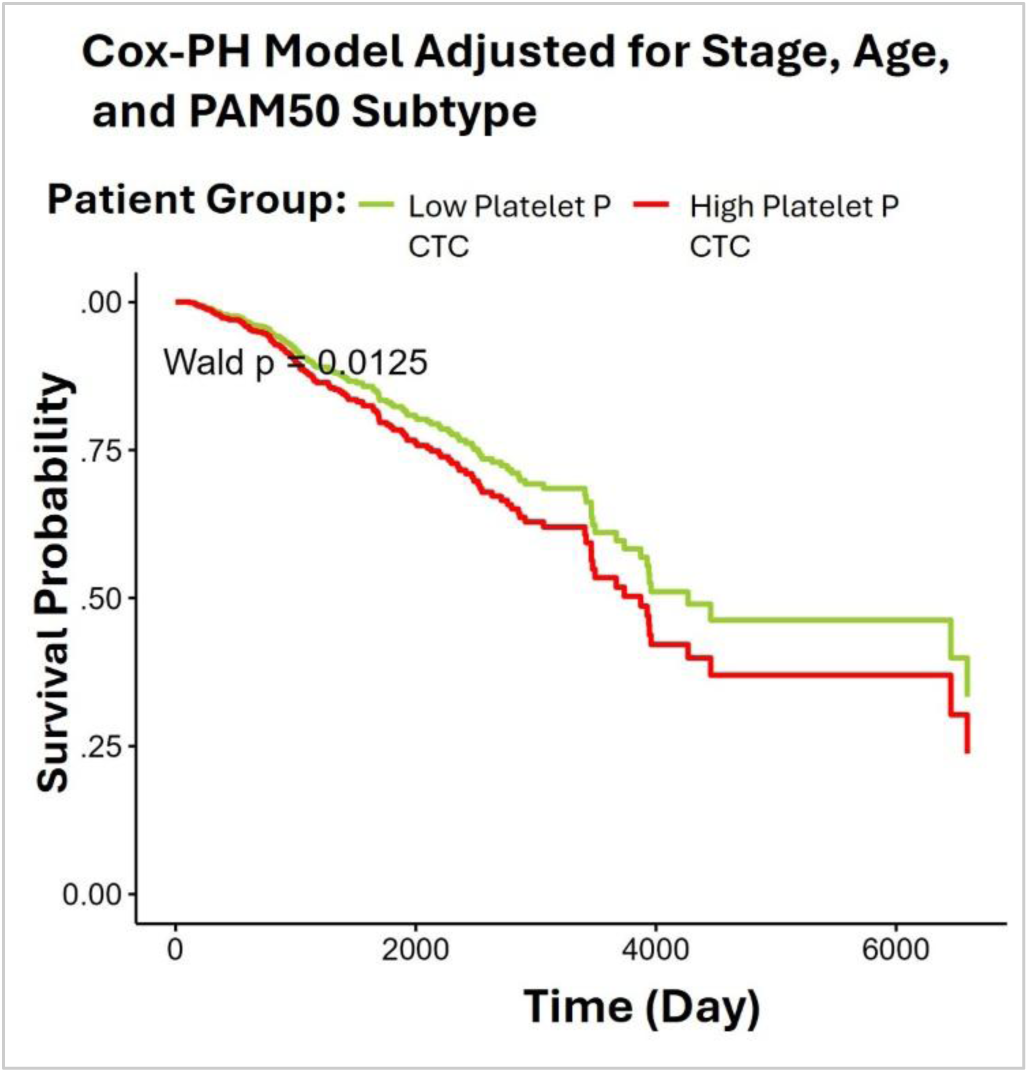
Cox-PH Model survival curves. Adjusted survival curves from a Cox-PH model comparing patients with high versus low estimated platelet-programmed tumor cell proportion, dichotomized at the cohort median. Curves represent marginal (average) adjusted survival probabilities. The p-value corresponds to the Cox model effect of platelet-programmed tumor cell proportion group. The model includes platelet-programmed tumor cell proportion group, adjusting for age, stage, PAM50.Platelet-P CTC: Platelet-Programmed CTC.

### 3.5 Communication between Platelet-programmed CTCs and Monocytes

To characterize intercellular signaling between Platelet-P CTCs and monocytes, ligand– receptor interaction analysis was performed with Platelet-P CTCs designated as signal-sending cells and onocytes as signal-receiving cells. The top-ranked ligand– receptor pairs are visualized in Fig.5, where ligands are displayed in the left semicircles and their corresponding receptors in the right semicircles. The color intensity of each semicircle represents the expression level of the respective ligand or receptor (Fig.5). The semicircle size represents the percentage of ligand/receptor expression in the corresponding cell type. Notably, the predicted pairs included several established myeloid immune checkpoints—HLA-G– LILRB1, HLA-G–LILRB2 and CD47–SIRPA—indicating that the analysis recovers interactions with independently documented checkpoint function and supporting the validity of the prediction pipeline. A complete list of identified ligand–receptor pairs is provided in Supplementary File 2. The ligand activity was estimated using the area under the precision-recall curve (AUPR) (Supplementary Fig.4).

**Fig 5.**
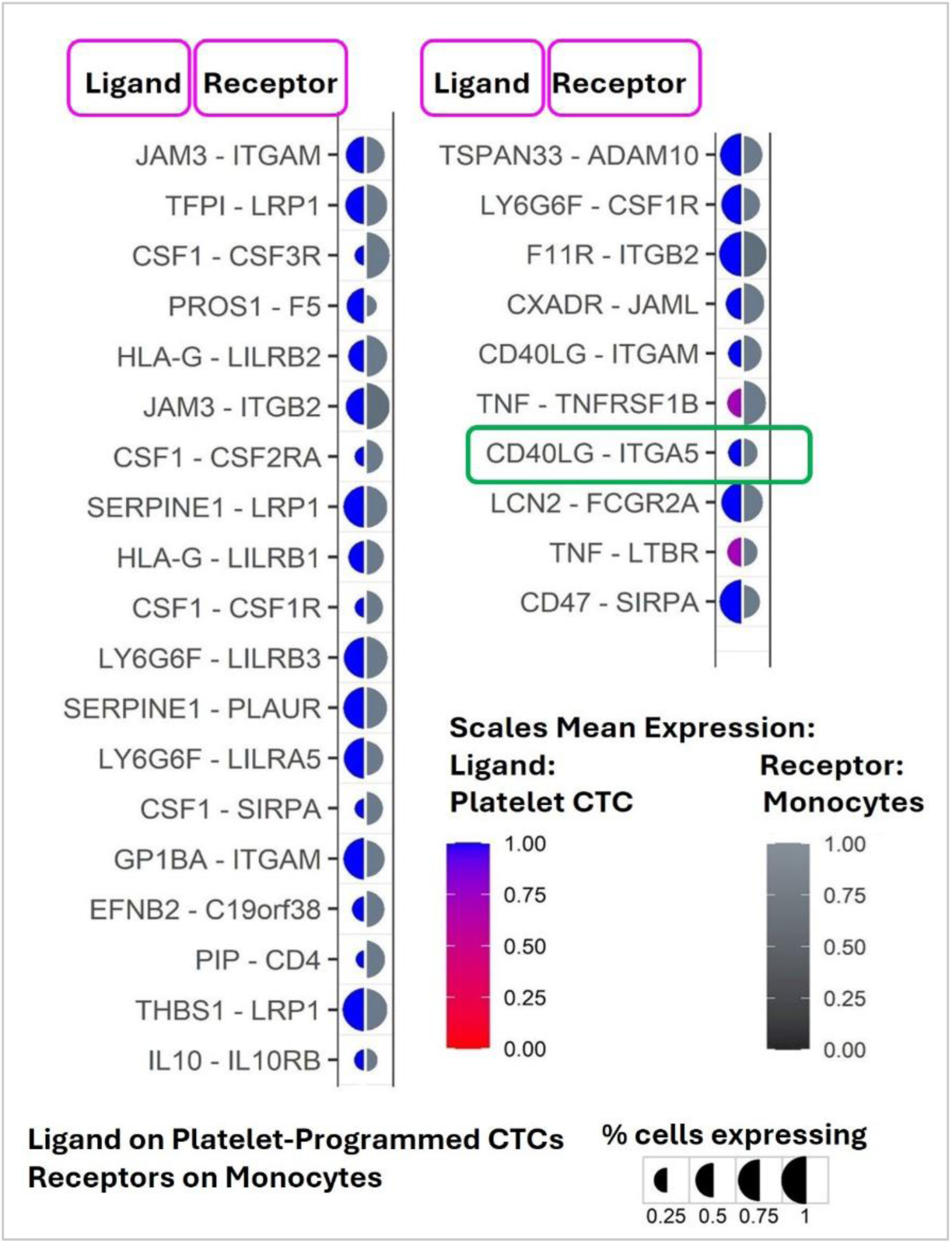
The predicted ligand-receptor pairs. Ligands (semicircles on the left) were detected on Platelet-Programmed CTCs and receptors (semicircles on the right) on Monocytes. The area of semicircles represents the percentage of ligand/receptor pairs on specific cell types. Platelet-P CTC, Platelet-Programmed CTC.

### 3.6 CD40LG–ITGA5:ITGB1 is a plausible “myeloid checkpoint–like” axis

The predicted CD40LG–ITGA5 ligand–receptor pair (highlighted in green, Fig.5) was selected against two criteria applied in sequence. First, both partners had to be expressed in approximately half of the cells of their respective populations, with ITGA5 detected in approximately 44% of monocytes. Second, and decisively, CD40LG–ITGA5 was the only predicted pair to meet the interaction-energy thresholds in both docking platforms (Fig.6). This pair was predicted at the transcriptional level, from ligand activity and receptor expression. To determine whether the predicted pair is also structurally capable of forming a stable complex, we assessed the interaction at the protein level using two rigid-body docking algorithms. Both returned interaction energies beyond the thresholds set by the experimentally resolved TNFα–TNFR reference complex (Hex score<−515; PyDock score<−45), indicating that the two proteins are structurally compatible with stable binding (Fig.6). The detailed settings and parameters of the Hex docking process are written in Supplementary Table.2. To define a threshold for near-native binding affinity, the experimentally resolved TNFα–TNFR complex (PDB: 7KP8), whose structure was determined by X-ray crystallography, was first docked on both platforms as a reference standard (Supplementary Table.3).

**Fig 6.**
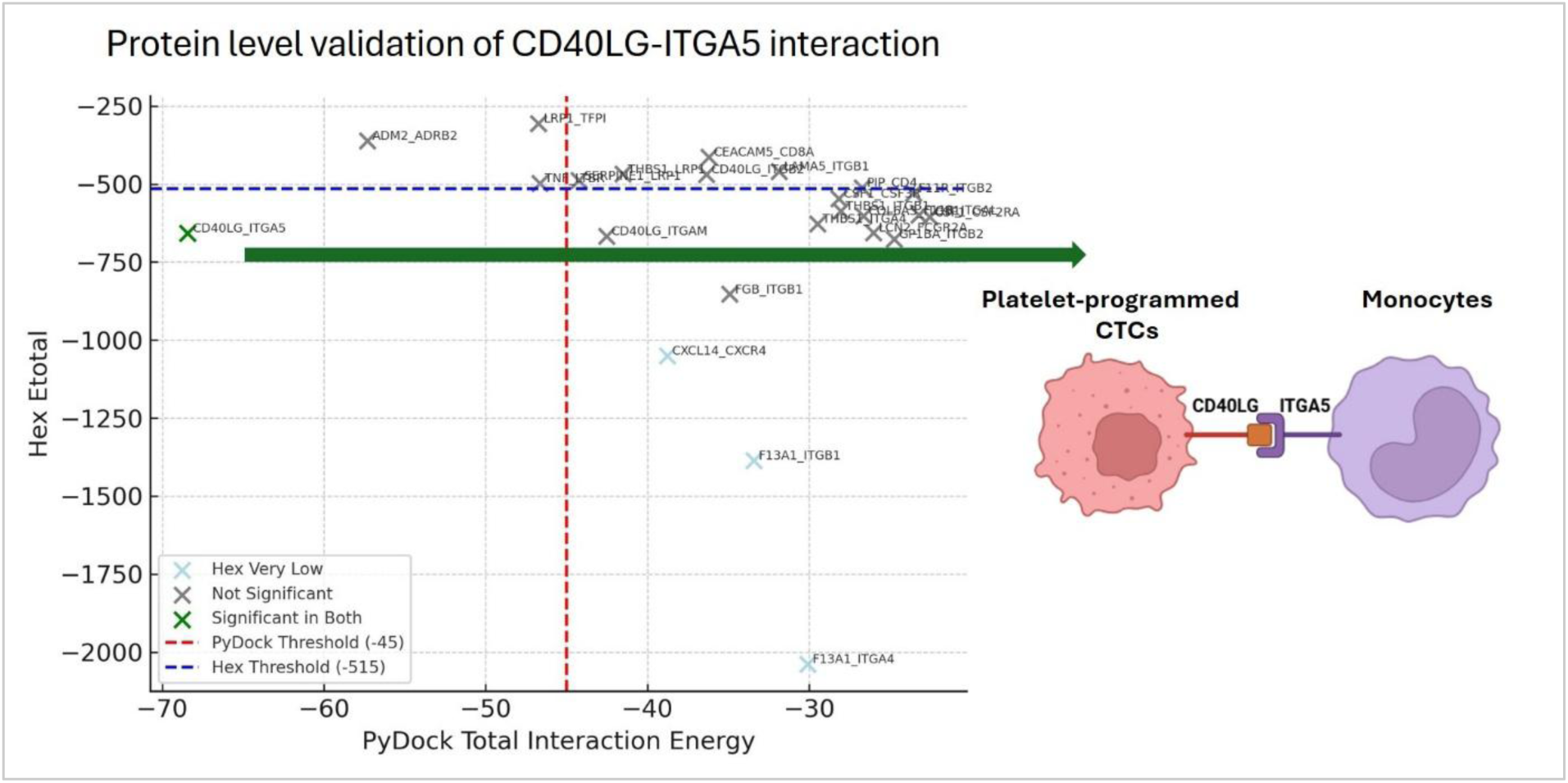
Protein-level structural assessment of the predicted ligand–receptor pair. Interaction energies below −45 for PyDock and below −515 for Hex were considered indicative of significant binding.

A targeted literature review of all predicted ligand–receptor pairs identified CD40LG–ITGA5 as the only pair not previously characterized as an immune checkpoint interaction, whereas several other predicted pairs had prior reported roles in immune regulation (Supplementary File 3). Supporting the biological relevance of this interaction, ITGB1—the obligatory binding partner of ITGA5 in the integrin heterodimer—was broadly expressed across leukocytes, including the majority of monocytes (Fig.7a). Assessment of the ITGA5:ITGB1 downstream signalling machinery (Section 2.8) showed that monocytes expressed each tier of the cascade: proximal adhesion components (PTK2B, TLN1, VCL), NF-κB subunits spanning canonical and non-canonical arms (REL, NFKB2, RELB), and cytokine, cytokine-receptor and TAM-associated signalling genes (IL10RA, CSF1R, CXCL8, HAVCR2, TGFBR1, TGFBR2, TGFBI, S100A9, VEGFA) (Fig.7b). Expression of this panel was largely restricted to the monocyte cluster relative to the T-cell and NK/NK-T clusters, indicating that the signalling apparatus for this axis is present, and preferentially so, in the receiver population (Supplementary File 3, Supplementary Table.4). Whether these programmes are actively engaged by Platelet-P CTCs was not tested here and would require comparison of monocyte states between patients differing in Platelet-P CTC burden.

**Fig 7.**
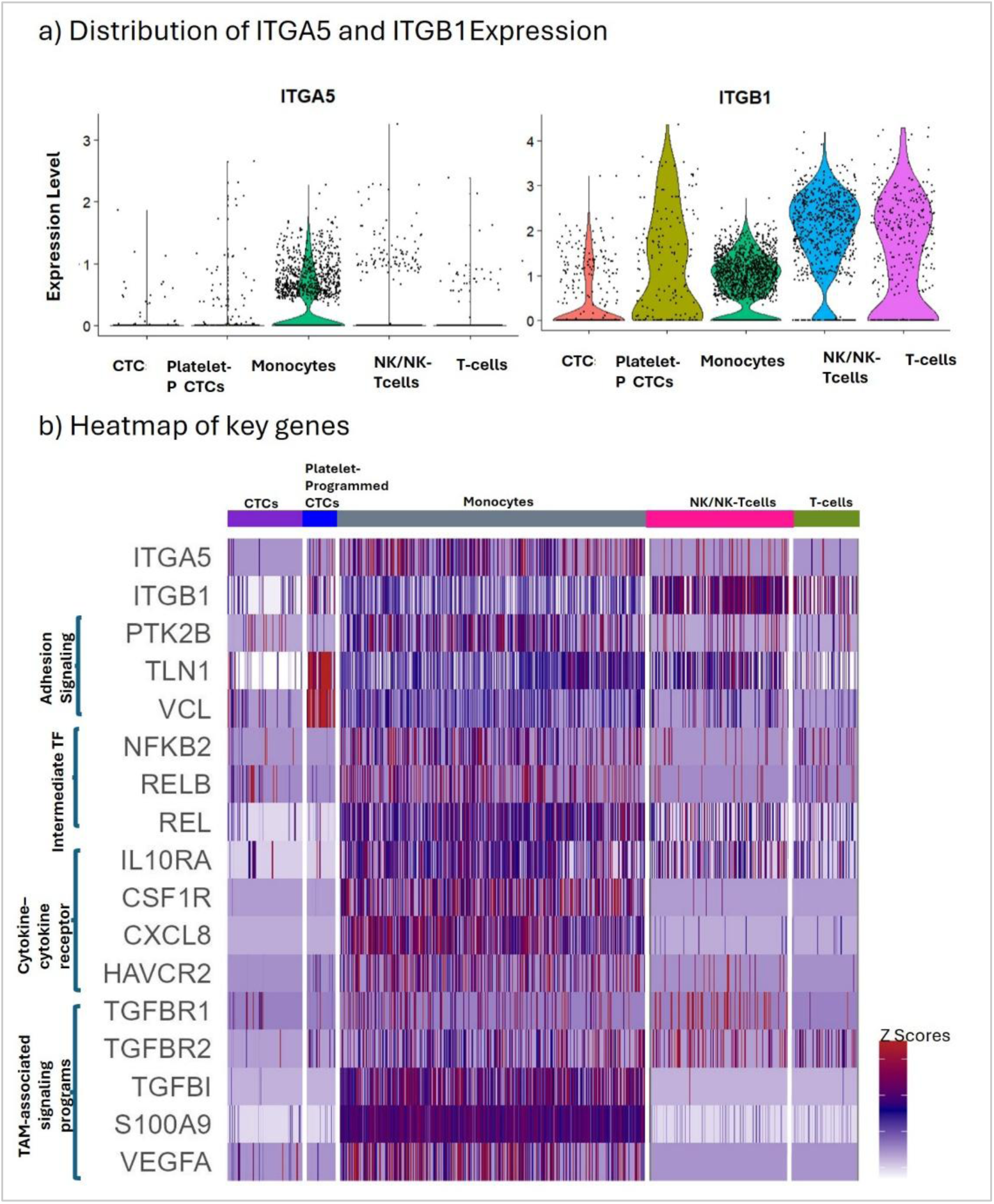
Expression of ITGA5:ITGB1 and its downstream signalling effectors across CTC and leukocyte clusters. A) Violin plot of ITGA5 and ITGB1 b) Heatmap of important genes. Platelet-P CTC: Platelet-programmed CTC

## 4 DISCUSSION

Metastatic dissemination of cancer cells from the primary tumor to distant sites remains a major cause of cancer-related mortality [1]. A critical step in this process is the survival of CTC subpopulations within the bloodstream, particularly those exhibiting platelet-associated transcriptional features [1, 3]. Nevertheless, immune surveillance of such CTC subpopulations in the bloodstream—a pivotal phase of metastasis—remains poorly defined. Therefore, novel immune checkpoint candidates targeting them warrant further exploration and validation.

In this study, integrative transcriptomic analyses suggest that Platelet-P CTCs may contribute to immune evasion and adverse clinical outcomes in breast cancer. A higher platelet-programmed tumor cell proportion was associated with increased mortality risk, supporting its potential prognostic significance. Furthermore, intercellular communication modeling identified CD40LG– ITGA5 as a candidate immunoregulatory interaction between Platelet-P CTCs and monocytes.

Computational docking analyses supported the structural plausibility of this interaction. Together, these findings raise the possibility that CD40LG–ITGA5 signaling may represent a previously unrecognized myeloid checkpoint-like axis in the circulation, although functional validation is required.

CTCs rapidly interact with circulating platelets, which become activated and form protective aggregates around tumor cells [1]. This platelet coating not only shields CTCs from immune-mediated clearance but also promotes metastatic traits, including epithelial–mesenchymal transition, angiogenesis, and extravasation [31]. CD40LG, a member of the tumor necrosis factor (TNF) superfamily, is a potent mediator of inflammatory and immune signaling. Upon release from activated platelets, CD40LG can stimulate endothelial cells to generate reactive oxygen species and upregulate adhesion molecules and chemokines, thereby enhancing vascular adhesion and facilitating metastatic dissemination [32, 33]. CD40LG has also been shown to promote angiogenesis through increased VEGF expression [34].

In our analysis, CD40LG expression was detected in a substantial fraction of Platelet-P CTCs, supporting the possibility that these cells retain platelet-derived immunomodulatory traits in circulation. Although CD40 represents the canonical receptor for CD40LG, it was not significantly expressed in monocyte clusters in our dataset. This observation aligns with previous studies indicating that CD40 expression is more prominent in T cells and neutrophils than in monocytes [32, 33]. Together, these findings raise the possibility that CD40LG expressed on Platelet-P CTCs may engage non-canonical receptors in monocytes, thereby contributing to immune modulation in the bloodstream. Importantly, while prior studies have primarily focused on CD40–CD40LG interactions in primary tumors, our work provides novel evidence that CD40LG is expressed on Platelet-P CTCs and may interact with non-canonical receptors in monocytes. This suggests a new immune checkpoint axis, not previously explored at the single-cell level in CTC subpopulations. However, it is important to acknowledge that the limited number of cells in the Platelet-Programmed CTC cluster represents a limitation of our study and warrants further validation in larger cohorts.

ITGA5 was expressed in approximately 44% of monocytes in our dataset, indicating that a substantial subset of monocytes may be capable of engaging CD40LG-mediated signaling. This expression frequency supports the biological plausibility of ITGA5 functioning as a receptor candidate in this context. Integrins like ITGA5 mediate cell–ECM adhesion and influence survival, migration, and proliferation [35, 36]. ITGA5-mediated signaling also contributes to cancer cell resistance to apoptosis [35, 37]. Prior work by Takada et al. shows that soluble CD40LG binds integrin heterodimers such as ITGA5:ITGB1, ITGAV:ITGB3, and ITGA4:ITGB1, all involved in CD40 signaling [38, 39]. They further reported that the ITGA5:ITGB1 (α5β1) integrin complex can function as a receptor for IL1B [40], underscoring the capacity of integrin heterodimers to mediate non-canonical immunoregulatory signaling. In our dataset, ITGB1 was co-expressed with ITGA5 in monocytes, supporting the potential formation of integrin α5β1 complexes. Although ITGB1 was not independently predicted as a receptor in our ligand–receptor modeling, integrins function as αβ heterodimers rather than as individual subunits. Therefore, it is biologically plausible that CD40LG may engage ITGA5 in the context of ITGA5:ITGB1 complexes on monocytes.

Monocytes and macrophages frequently adopt pro-tumoral phenotypes within the tumor microenvironment, contributing to immune suppression [41]. ITGA5 has been implicated in these processes. For example, Pantano et al. demonstrated that ITGA5 promotes bone metastasis in breast cancer by enhancing extracellular matrix adhesion [42], while White et al. reported that integrin α5β1 (ITGA5:ITGB1) expression on monocytes induces CXC chemokines with pro-angiogenic properties [43]. Although these studies did not directly examine monocyte–CTC interactions, they suggest that ITGA5-mediated signaling may enhance pro-tumoral and immunomodulatory functions [31, 42–44]. In this context, ITGA5⁺ monocytes may participate in ligand–receptor interactions that influence immune cell activation states.

Takada et al. further demonstrated that CD40LG must bind integrins like ITGA5:ITGB1 to fully activate CD40-mediated immune signaling, including NF-κB and cytokine production [39]. While this work did not specifically focus on cancer, it highlights a vital immune mechanism relevant to tumor immunity and inflammation. The significance of this signaling axis is further contextualized by Nirschl et al., who demonstrated that mononuclear phagocytes—including monocytes, macrophages, and dendritic cells—share a conserved homeostatic gene program centered on NF-κB signaling, notably involving the non-canonical subunit RelB, that is co-opted across multiple human cancers to suppress anti-tumor immunity [45]. Strikingly, CD40 was identified as one of the transcripts within this program, and the authors proposed that its activity within the myeloid compartment may confer greater CD40 and IL-12 dependence in the tumor microenvironment [45]. The conservation of this program across monocytes and macrophages — not only dendritic cells — is particularly relevant, as it suggests that CD40LG–ITGA5:ITGB1-driven NF-κB activation in monocytes may similarly engage or be co-opted by this homeostatic axis. Accordingly, investigating CD40LG–ITGA5:ITGB1 interactions in tumor-associated monocytes could guide the development of cancer immunotherapies targeting myeloid checkpoints, particularly by disrupting the tolerogenic programming that tumors exploit to evade immune surveillance.

The curated ITGA5:ITGB1 effector panel was expressed across monocytes, spanning adhesion, NF-κB and immunoregulatory tiers, including IL10RA, TGFBR1/2, CSF1R, HAVCR2, S100A9 and VEGFA. Several of these genes are constitutively expressed in circulating monocytes, so their presence indicates capacity rather than activation. Nonetheless, co-expression of the complete cascade—receptor, proximal adaptors, transcription factors and downstream immunoregulatory effectors—within the same population indicates that monocytes are equipped to transduce CD40LG–ITGA5:ITGB1 signalling. Whether engagement of this axis shifts monocytes toward an immunosuppressive, TAM-like state remains to be tested.

Finally, a limitation of this study is that it relied on bioinformatics and computational biology methods. While we attempted to integrate gene expression data from various sources and assess the ligand-receptor interactions through molecular docking, the results should be interpreted with caution. Confirmatory experiments and clinical validation are essential to substantiate these findings and translate them into effective medical interventions.

## 5 CONCLUSION

Our findings reveal a novel CD40LG–ITGA5 interaction between Platelet-P CTCs and monocytes, suggesting a potential immune checkpoint axis in cancer. This interaction may contribute to tumor immune evasion and metastasis, highlighting its promise as a target for future immunotherapeutic strategies. Further experimental validation is needed to confirm its clinical relevance.

## Data Availability Statement

Publicly available datasets were analyzed in this study. Single-cell RNA-seq data are available from NCBI GEO under the accession numbers listed in Table 1; TCGA-BRCA bulk RNA-seq and clinical data were obtained from the GDC Data Portal (https://portal.gdc.cancer.gov/). Processed results are provided in the Supplementary Material.

## Ethics Statement

Ethical review and approval was not required for this study because only publicly available, de-identified datasets were analyzed.

## Author Contributions

S.K. contributed significantly to the study’s conceptualization, formal analysis, methodology, and original draft, and also participated in the manuscript revision, editing, and supervision. H.B. Docking analysis and preparing docking results. M.Z. Data curation and investigation, visualization, and preparing results. Z.S. Data collection. Y.S. Data collection. N.H. Visualization and original drafting of part of the introduction. M.S.F. Methodology, review, and editing. N.A. Review and editing.

All authors reviewed and approved the final manuscript.

## Funding

The author(s) received no financial support for the research, authorship, and/or publication of this article.

## Conflict of Interest

The authors declare that the research was conducted in the absence of any commercial or financial relationships that could be construed as a potential conflict of interest.

## Acknowledgments

No substantial contributions were received from individuals who do not meet the criteria for authorship.

## Abbreviations

Platelet-P CTCs: Platelet-programmed CTCs are circulating tumor cells that retain platelet-associated transcriptional programs consistent with prior platelet-mediated signaling, without evidence of persistent physical platelet association

## Notes

### Competing Interest Statement

The authors have declared no competing interest.

## REFERENCES

1. Liu, X., et al., Immune checkpoint HLA-E: CD94-NKG2A mediates evasion of circulating tumor cells from NK cell surveillance. Cancer Cell, 2023. 41(2): p. 272–287. e9.

2. Dai, C.S., et al., Circulating tumor cells: Blood-based detection, molecular biology, and clinical applications. Cancer Cell, 2025. 43(8): p. 1399–1422.

3. Gautam, D., et al., Platelets and circulating (tumor) cells: partners in promoting metastatic cancer. Current Opinion in Hematology, 2025. 32(1): p. 52–60.

4. He, X. and C. Xu, Immune checkpoint signaling and cancer immunotherapy. Cell research, 2020. 30(8): p. 660–669.

5. Nagano, M., et al., PD-L1 expression on circulating monocytes in patients with breast cancer. Annals of Oncology, 2018. 29: p. ix10.

6. Zhao, J., et al., The MHC class I-LILRB1 signalling axis as a promising target in cancer therapy. Scandinavian journal of immunology, 2019. 90(5): p. e12804.

7. Dumont, C., et al., CD8+ PD-1–ILT2+ T cells are an intratumoral cytotoxic population selectively inhibited by the immune-checkpoint HLA-G. Cancer immunology research, 2019. 7(10): p. 1619–1632.

8. Tang, J., et al., The clinical trial landscape for PD1/PDL1 immune checkpoint inhibitors. Nature reviews Drug discovery, 2018. 17(12): p. 854–855.

9. Lin, A. and W.-H. Yan, HLA-G/ILTs targeted solid cancer immunotherapy: opportunities and challenges. Frontiers in Immunology, 2021. 12: p. 698677.

10. Jordan, N.V., et al., HER2 expression identifies dynamic functional states within circulating breast cancer cells. Nature, 2016. 537(7618): p. 102–106.

11. Szczerba, B.M., et al., Neutrophils escort circulating tumour cells to enable cell cycle progression. Nature, 2019. 566(7745): p. 553–557.

12. Gkountela, S., et al., Circulating tumor cell clustering shapes DNA methylation to enable metastasis seeding. Cell, 2019. 176(1): p. 98–112. e14.

13. Ebright, R.Y., et al., Deregulation of ribosomal protein expression and translation promotes breast cancer metastasis. Science, 2020. 367(6485): p. 1468–1473.

14. Diamantopoulou, Z., et al., The metastatic spread of breast cancer accelerates during sleep. Nature, 2022. 607(7917): p. 156–162.

15. Poonia, S., et al., Marker-free characterization of full-length transcriptomes of single live circulating tumor cells. Genome Research, 2023. 33(1): p. 80–95.

16. Young, M.D. and S. Behjati, SoupX removes ambient RNA contamination from droplet-based single-cell RNA sequencing data. Gigascience, 2020. 9(12): p. giaa151.

17. Pauken, C.M., et al., Heterogeneity of circulating tumor cell neoplastic subpopulations outlined by single-cell transcriptomics. Cancers, 2021. 13(19): p. 4885.

18. Butler, A., et al., Integrating single-cell transcriptomic data across different conditions, technologies, and species. Nature biotechnology, 2018. 36(5): p. 411–420.

19. Becht, E., et al., Dimensionality reduction for visualizing single-cell data using UMAP. Nature biotechnology, 2019. 37(1): p. 38–44.

20. Kuleshov, M.V., et al., Enrichr: a comprehensive gene set enrichment analysis web server 2016 update. Nucleic acids research, 2016. 44(W1): p. W90–W97.

21. Schmiedel, B.J., et al., Impact of genetic polymorphisms on human immune cell gene expression. Cell, 2018. 175(6): p. 1701–1715. e16.

22. Chen, L., et al., Genetic drivers of epigenetic and transcriptional variation in human immune cells. Cell, 2016. 167(5): p. 1398–1414. e24.

23. Mabbott, N.A., et al., An expression atlas of human primary cells: inference of gene function from coexpression networks. BMC genomics, 2013. 14(1): p. 632.

24. Street, K., et al., Slingshot: cell lineage and pseudotime inference for single-cell transcriptomics. BMC genomics, 2018. 19(1): p. 477.

25. Browaeys, R., W. Saelens, and Y. Saeys, NicheNet: modeling intercellular communication by linking ligands to target genes. Nature methods, 2020. 17(2): p. 159–162.

26. Tsoucas, D., et al., Accurate estimation of cell-type composition from gene expression data. Nature communications, 2019. 10(1): p. 2975.

27. Finak, G., et al., MAST: a flexible statistical framework for assessing transcriptional changes and characterizing heterogeneity in single-cell RNA sequencing data. Genome biology, 2015. 16: p. 1–13.

28. Therneau, T.M., Survival Analysis [R package survival version 2.42-6]. 2015.

29. Guex, N. and M.C. Peitsch, *SWISS-MODEL and the Swiss-Pdb Viewer: an environment for* comparative *protein modeling*. electrophoresis, 1997. 18(15): p. 2714–2723.

30. Jiménez-García, B., C. Pons, and J. Fernández-Recio, pyDockWEB: a web server for rigid-body protein–protein docking using electrostatics and desolvation scoring. Bioinformatics, 2013. 29(13): p. 1698–1699.

31. Liu, Y., et al., Platelet-mediated tumor metastasis mechanism and the role of cell adhesion molecules. Critical Reviews in Oncology/Hematology, 2021. 167: p. 103502.

32. Cognasse, F., et al., Platelets as key factors in inflammation: focus on CD40L/CD40. Frontiers in immunology, 2022. 13: p. 825892.

33. Ara, A., K.A. Ahmed, and J. Xiang, Multiple effects of CD40–CD40L axis in immunity against infection and cancer. ImmunoTargets and therapy, 2018: p. 55–61.

34. Pazoki, A., et al., Soluble CD40 Ligand as a Promising Biomarker in Cancer Diagnosis. Cells, 2024. 13(15): p. 1267.

35. Hou, J., et al., The roles of integrin α5β1 in human cancer. OncoTargets and therapy, 2020: p. 13329–13344.

36. Liu, F., et al., Integrins in cancer: Emerging mechanisms and therapeutic opportunities. Pharmacology & therapeutics, 2023. 247: p. 108458.

37. Yang, Y., et al., Integrin α5 promotes migration and invasion through the FAK/STAT3/AKT signaling pathway in icotinib-resistant non-small cell lung cancer cells. Oncology Letters, 2021. 22(1): p. 1–11.

38. Takada, Y.K., et al., Soluble CD40L activates soluble and cell-surface integrin αvβ3, α5β1, and α4β1 by binding to the allosteric ligand-binding site (site 2). Journal of Biological Chemistry, 2021. 296.

39. Takada, Y.K., et al., Integrin binding to the trimeric interface of CD40L plays a critical role in CD40/CD40L signaling. The Journal of Immunology, 2019. 203(5): p. 1383–1391.

40. Takada, Y.K., et al., Direct binding to integrins and loss of disulfide linkage in interleukin-1β (IL-1β) are involved in the agonistic action of IL-1β. Journal of Biological Chemistry, 2017. 292(49): p. 20067–20075.

41. Fendl, B., et al., Macrophage and monocyte subsets as new therapeutic targets in cancer immunotherapy. ESMO open, 2023. 8(1): p. 100776.

42. Pantano, F., et al., Integrin alpha5 in human breast cancer is a mediator of bone metastasis and a therapeutic target for the treatment of osteolytic lesions. Oncogene, 2021. 40(7): p. 1284–1299.

43. White, E.S., et al., Monocyte-fibronectin interactions, via α5β1 integrin, induce expression of CXC chemokine-dependent angiogenic activity. The Journal of Immunology, 2001. 167(9): p. 5362–5366.

44. Zhang, Q., et al., The interplay between integrins and immune cells as a regulator in cancer immunology. International Journal of Molecular Sciences, 2023. 24(7): p. 6170.

45. Nirschl, C.J., et al., IFNγ-dependent tissue-immune homeostasis is co-opted in the tumor microenvironment. Cell, 2017. 170(1): p. 127–141. e15.

