## Supplemental Materials for "Platelet-programmed circulating tumor cells signal to monocytes through a candidate CD40LG–ITGA5:ITGB1 myeloid checkpoint axis in breast cancer"

Khoshbakht S et al.

Supplementary:

### 1 Data:

After integrating different CTC and WBC data downloaded from NCBI, we had 706 CTCs and 2925 WBCs before QC filtering (Fig.S1). WBCs with low quality were filtered out. CTCs with the absence of at least one of the CTC markers, including KRT18, KRT19, KRT8, EPCAM, or KRT7, were filtered out. Moreover, CTCs with PTPRC-positive cells were removed from downstream analyses(Suplementary Fig.2).

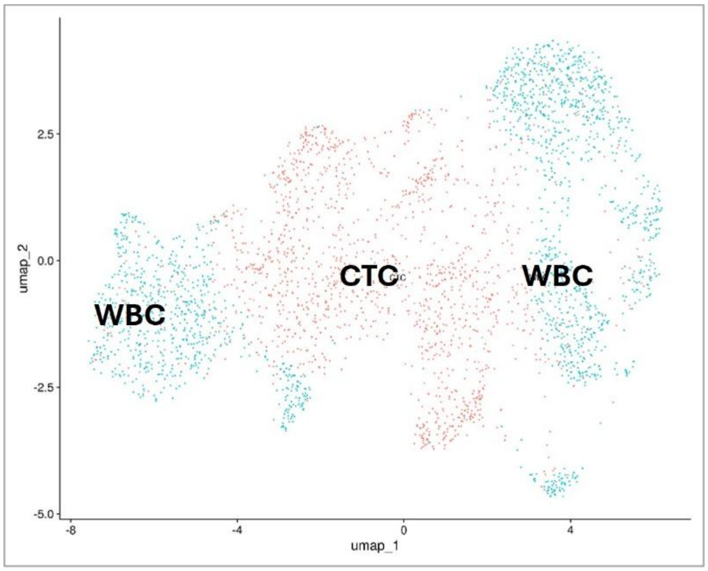


*Supplementary Fig.1 CTCs and WBCs DimPlot.*
*Red color dots indicate CTC, and turquoise dots indicate the WBCs.*

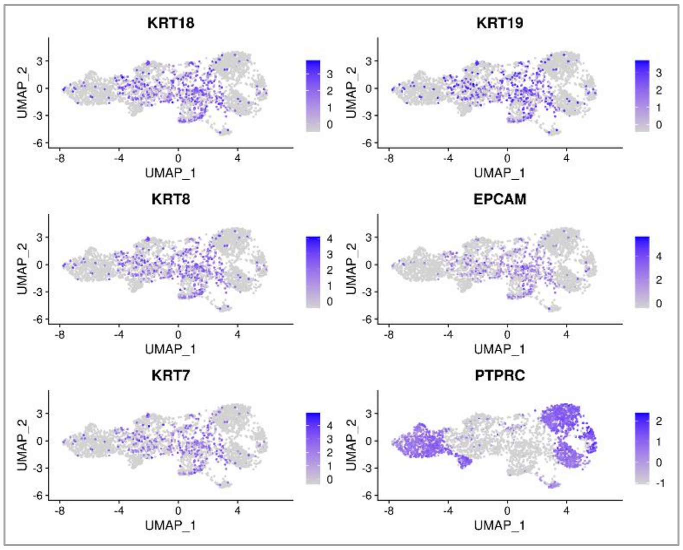


*Supplementary Fig.2 Feature Plot of marker genes*

### 2 Annotation of subpopulation:

The marker genes utilized for the annotation of platelet-Programmed CTCs (CTCs with platelet markers), T cells, Monocytes, CTCs, and NK/NK-Tcells were reported in Supplementary Table.1. Most Platelet-like marker genes are cell surface markers. Moreover, we implemented 4 computational methods to annotate the subpopulation of CTCs and WBCs. Supplementary Fig.3a,b, and C indicate the WBC annotation, and d indicates the CTC subpopulation annotation (Supplementary Fig.3a,b,c, and d).

*Supplementary Table.1: Marker genes*

| Platelet-like markers | T-cells | Monocytes | NK/NK-T-cells | CTC |
|---|---|---|---|---|
| PTPRC | CD3D | FGFBP2 | CD3D | EPCAM |
| PECAM1 | CD3E | PRF1 | IL7R | KRT8 |
| GP1BB | IL7R | FCGR3A | LEF1 | KRT18 |
| GP5 | LEF1 | KLRD1 | CD3E | KRT19 |
| SELP | CD8A | FCGR3A | KIR2DL3 | ERBB2 |
| ITGB3 | CD8B | CTSW | KIR2DL1 | JUP |
| CD97 | | GZMA | GZMB | YAP1 |
| SELPLG | | GZMB | PRF1 | ICAM1 |
| TGFB1 | | GZMH | CD97 | IL8 |
| TNFSF18 | | CD3E | | VEGFA |
| YAP1 | | | | NFE2L2 |
| KLRK1 | | | | VIM |
| PDGFB | | | | |
| ITGA2B | | | | |
| PPBP | | | | |
| RGS18 | | | | |
| PF4 | | | | |
| GP9 | | | | |
| GP1BA | | | | |

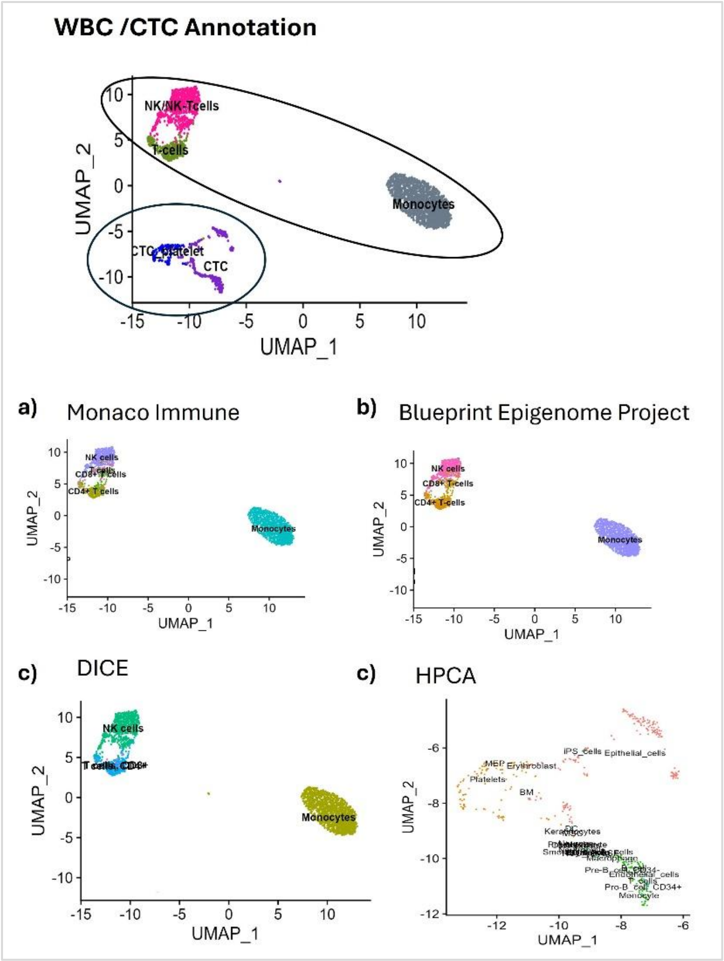


*Supplementary Fig.3. Annotation of subpopulation*

### 3 Predicted ligand activities:

Based on the NicheNet method, a ligand-target pair receives a high regulatory potential if the regulators of the prior target gene are downstream of the signaling network of the ligand. This procedure, called ligand activity prediction, ranks ligands according to how well their prior target gene predictions correspond to the observed gene expression changes resulting from communication with sender cells. The personalized Page Rank (PPR) method was used for ligand-target potential scores. In PPR, the ligand of interest gets the value of 1, and the other genes get 0. The ligand activity in this work was calculated using the area under the precision-recall curve (AUPR) (Supplemenatry Fig.4). AUPR shows the target gene prediction ability.

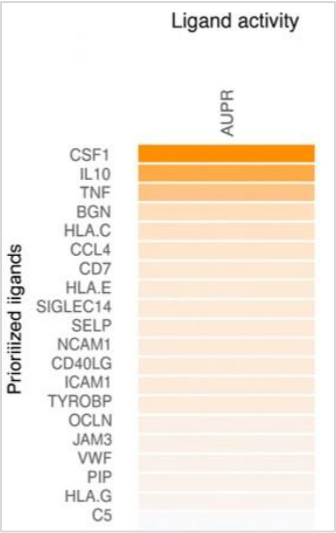

*Supplementary Fig.4 Ligand activity*

### 4 Molecular Docking

To estimate the interaction energy of the predicted ligand-receptor pairs at the protein level, we implemented two types of molecular docking. Estimating a probable stable interaction between predicted proteins requires an appropriate evaluation of the intermolecular energetics. Since protein-protein docking algorithms often do not utilize flexible docking approaches due to the high number of residues for calculation, electrostatics and VdW interactions become crucial for calculations. Each algorithm possesses a unique scoring method for prioritization of the docking positions. However, all scores are relative and must be compared with an original or naturally existing source. To overcome this weakness, we carried out docking with an online webserver (PyDock) and an offline software (Hex).

The detailed settings and parameters of the Hex docking process are written in Supplementary Table.2.

*Supplementary Table.2 Settings and parameters of the Hex*

| Parameter | Value | Parameter | Value |
|---|---|---|---|
| Correlation type | Shape + Electro + DRAS | Ligand Range | 180 |
| FFT Mode | 3D | Twist Range | 360 |
| Sampling Method | Range Angels | Distance Range | 40 |
| Post Processing | OPLS Energies | Translation Step | 0.8 |
| Steric Scan | 18 | Receptor Range | 180 |
| Final Search | 30 | Solutions | 2000 |
| Grid Dimensions | 0.6 | Box Size | 10 |

The natural/near-natural binding positions of the TNFα-TNFR complex in both platforms were mentioned (Supplementary Table.3).

*Supplementary Table.3 TNFα-TNFR binding positions*

| **PyDock** | | | | | **Hex** | | | |
|---|---|---|---|---|---|---|---|---|
| **Electrostatics** | **Desolvation** | **VdW** | **ETotal** | **Ranking** | **Etotal** | **Eshape** | **Eforc** | **RMS** |

| | | | | | | | e | |
|---|---|---|---|---|---|---|---|---|
| -22.678 | -41.244 | 24.18 | -61.504 | 1 | -1398.38 | -1679.82 | 281.44 | 0.32 |
| -21.871 | -40.01 | 46.973 | -57.184 | 2 | -645.09 | -1111.03 | 465.94 | 27.45 |
| -38.016 | -18.339 | 10.094 | -55.345 | 3 | -557.73 | -675.24 | 117.51 | 39.47 |
| -37.033 | -13.938 | -6.308 | -51.602 | 4 | -534.47 | -748.57 | 214.1 | 39.11 |
| -33.582 | -16.438 | 21.463 | -47.873 | 5 | -517.46 | -613.9 | 96.43 | 39.14 |

Supplementary Table.4 includes the docking energies of the best poses of each protein-protein docking pair. Compared with interaction energies (IEs) of the TNFα-TNFR complex, if the IEs for PyDock and Hex drop lower than -45 and -515, the probability of natural interaction also increases.

According to supplementary Table.4, the protein pairs with yellow-highlighted IEs could create a relatively stable interaction. However, the number of proteins that are recognized as potentially interactable in both PyDock and Hex platforms is low and highlighted in green. Supplementary Table.4 includes the docking energies of the best poses of each protein-protein docking pair.

*supplementary Table 4 Docking Energies for Protein pairs in Monocytes*

| Cluster | PyDock | | | | | Hex | | | |
|---|---|---|---|---|---|---|---|---|---|
| | **Protein Pairs** | **Electrostatics** | **Desolvation** | **VdW** | **Total** | **Etotal** | **Eshape** | **Eforce** | **RMS** |
| Monocytes | LRP1_TFPI | -25.586 | -27.304 | 61.607 | -46.73 | -305.5 | -360.47 | 54.97 | -1 |
| | CSF1_CSF3R | -35.415 | 5.592 | 16.557 | -28.167 | -467.97 | -546.54 | 78.57 | -1 |
| | CSF1_CSF2RA | -19.607 | -4.332 | 13.88 | -22.551 | -606.07 | -683.6 | 77.53 | -1 |
| | SERPINE1_LRP1 | -43.281 | -1.251 | 3.091 | -44.223 | -488.8 | -566.96 | 78.16 | -1 |
| | PIP _CD4 | -28.437 | 2.524 | -8.107 | -26.724 | -511.63 | -694.09 | 182.47 | -1 |
| | THBS1_LRP1 | -51.612 | 7.328 | 27.594 | -41.524 | -468.19 | -529 | 60.81 | -1 |
| | CD40LG_ITGAM | -15.722 | -28.52 | 17.061 | -42.536 | -666.48 | -775.83 | 109.35 | -1 |
| | CD40LG_ITGA5 | -34.216 | -38.695 | 44.956 | -68.415 | -656.83 | -656.83 | 0 | -1 |
| | LCN2_FCGR2A | -31.221 | 9.826 | -46.168 | -26.012 | -656.07 | -908.66 | 252.59 | -1 |
| | TNF_LTBR | -26.006 | -23.314 | 26.854 | -46.635 | -497.64 | -593.39 | 95.75 | -1 |

*The yellow-colored cells indicate the significant pairs in Hex.*
*The green-colored cells indicate the significant pairs in Hex and PyDock.*
*Light blue cells indicate the significant pairs in Hex with very low energy.*
